# Chronic exercise protects against the progression of renal cyst growth and dysfunction in rats with polycystic kidney disease

**DOI:** 10.1101/2021.03.11.434857

**Authors:** Jiahe Qiu, Yoichi Sato, Lusi Xu, Takahiro Miura, Masahiro Kohzuki, Osamu Ito

**Author notes:** **Corresponding author:** Osamu Ito, MD, PhD, Division of General Medicine and Rehabilitation, Tohoku Medical and Pharmaceutical University, Faculty of Medicine, 1-15-1 Fukumoto, Miyagino-ku, Sendai 983-8536, Japan.

## Abstract

**Introduction:** Polycystic kidney disease (PKD) is a genetic disorder characterized by the progressive enlargement of renal epithelial cysts and renal dysfunction. Previous studies have reported the beneficial effects of chronic exercise on chronic kidney disease. However, the effects of chronic exercise have not been fully examined in PKD patients or models. The effects of chronic exercise on the progression of PKD were investigated in a polycystic kidney (PCK) rat model.

**Methods:** Six-week-old male PCK rats were divided into a sedentary group and an exercise group. The exercise group underwent forced treadmill exercise for 12 weeks (28 m/min, 60 min/day, 5 days/week). After 12 weeks, kidney function and histology were examined, protein expressions were analyzed, and signaling cascades of PKD were examined.

**Results:** Chronic exercise reduced the excretion of urinary protein, liver-type fatty acid-binding protein, plasma creatinine, urea nitrogen, and increased plasma irisin and urinary arginine vasopressin (AVP) excretion. Chronic exercise also slowed renal cyst growth, glomerular damage, and interstitial fibrosis, and led to reduced Ki-67 expression. Chronic exercise had no effect on cAMP content but decreased the renal expression of B-Raf and reduced the phosphorylation of extracellular signal-regulated kinase (ERK), mammalian target of rapamycin (mTOR), and S6.

**Conclusion:** Chronic exercise slows renal cyst growth and damage in PCK rats, despite increasing AVP, with down-regulation of the cAMP/B-Raf/ERK and mTOR/S6 pathways in the kidney of PCK rats.

## Introduction

Polycystic kidney disease (PKD) is the most prevalent of all genetic disorders, which is characterized by progressive enlargement of epithelial cysts in the kidney. Autosomal dominant PKD (ADPKD) is caused by mutations in *PKD1* (encoding polycystin-1) or *PKD2* (encoding polycystin-2), whereas autosomal recessive PKD (ARPKD) is caused by mutations in *PKHD1* (encoding fibrocystin). Polycystin-1, polycystin-2, and fibrocystin are all localized in the primary cilia and are required for the regulation of Ca^2+^ influx in response to ciliary bending. Primary cilia abnormalities are associated with lowered intracellular Ca^2+^ (1). Low intracellular Ca^2+^-related abnormal signaling leads to the induction of cyst epithelial cell proliferation, which is a key feature of cyst growth (2).

Low intracellular Ca^2+^ activates adenylyl cyclase and increases intracellular cAMP levels. Next, cAMP and protein kinase A signaling upregulates the B-Raf and extracellular signaling-regulated kinase (ERK) pathway in renal cyst epithelial cells (3). The finding that increased cAMP signaling is a crucial driver of cyst growth has led to the development of arginine vasopressin (AVP) type 2 receptor (V2R)-based therapy. Antagonists of V2R, including tolvaptan, reduce renal cAMP content by inhibiting V2 receptors which coupled with the stimulatory G protein (Gs) and slow cyst growth and the decline of renal function in ADPKD patients and rodent PKD models (4). In addition, it has been reported that the mammalian target of rapamycin (mTOR) and S6 pathway promotes cyst growth by enhancing the proliferation, size, and metabolism of renal tubular cells (5).

Lifestyle modifications that slow the progression of chronic kidney disease (CKD) have long been a topic of research interest. Clinical studies have reported that chronic exercise slows the decline in glomerular filtration rate (6, 7), decreases albuminuria (8), delays the initiation of dialysis, and diminishes overall mortality in CKD patients (9). We have also reported that chronic exercise at moderate intensity has renal protective effects in CKD model rats with 5/6 nephrectomy, diabetic nephropathy, and salt-sensitive hypertension (10-14). With respect to the mechanisms of the beneficial effects of chronic exercise, a newly discovered exercise-induced myokine, irisin, has been reported to have renal protective effects (15). Both endurance and resistance exercises increase irisin in skeletal muscles and plasma (16). Moreover, recombinant irisin administration prevents renal damage and fibrosis in mice with folic acid nephropathy, unilateral ureteral obstruction, and 5/6 nephrectomy (15).

ADPKD patients with a glomerular filtration rate ≥ 60 (mL/min/1.73 m^2^) have a low exercise capacity (17). Similarly, we recently reported a low exercise capacity in polycystic kidney (PCK) rats (18), which have polycystic kidney and liver diseases and resemble human ADPKD (19, 20). Chronic exercise at a moderate intensity for 12 weeks improved the low exercise capacity and, unexpectedly, slowed liver cyst growth and fibrosis in PCK rats (18). However, it is controversial whether chronic exercise has renal protective effects in PKD patients and/or models, because acute or chronic exercise stimulates the posterior pituitary gland to secrete AVP, thus increasing AVP levels (21). Therefore, we examined the effects of chronic exercise on the progression of PKD, as well as on the signaling cascades responsible for cellular proliferation in PCK rats.

## Methods

### Experimental animals

Five-week-old male PCK and Sprague-Dawley rats were obtained from Charles River Laboratories Japan (Yokohama, Japan). All rats had free access to tap water and were fed a standard rat diet (Labo MR Stock, Nosan Kogyo Co., Yokohama, Japan). All animal experiments were approved by the Tohoku University Committee for Animal Experiments and were performed in accordance with the Guidelines for Animal Experiments and Related Activities of Tohoku University (permit no. 2018-084).

### Exercise protocol

After 1 week of acclimatization, PCK rats were divided into the sedentary group (Sed-PCK, *n*=10) or the exercise group (Ex-PCK, *n*=10). The Sprague-Dawley (SD) rats were set as a control group (Con-SD, *n*=10). The Ex-PCK group underwent forced treadmill exercise with moderate intensity, using treadmills (KN-73; Natsume Industries, Tokyo, Japan) for 12 weeks with the following protocol: 28 m/min, 60 min/day, and 5 days/week (11).

### Plasma and urinary parameters

The rats were housed individually in metabolic cages (Model ST; Sugiyama-General, Tokyo, Japan) for 3 days to acclimatize to the conditions. Food and water intake were measured, and urine was collected on ice for 24h. Systolic blood pressure was measured using the tail-cuff method (MK-2000A; Muromachi Kikai, Tokyo, Japan). The rats were euthanized with sodium pentobarbitone (100 mg/kg, i.p.) and blood samples were collected from the ventral aorta. Urine and blood samples were centrifuged for 10 min at 2 000 × g, and the supernatant was collected. Plasma and urine aliquots were rapidly frozen and stored at –80°C until analysis.

Urinary protein and plasma glucose, total cholesterol, triglyceride, urea nitrogen, and creatinine were measured using standard auto-analysis techniques (SRL Inc., Tokyo, Japan). L-FABP was measured using a highly sensitive enzyme-linked immunosorbent assay (CMIC, Tokyo, Japan) (22). Plasma AVP levels are fluctuated by anesthetics or stress (23), and the indwelling catheter into the femoral artery may affect treadmill running. Therefore, we measured AVP concentration in the 24h urine by radioimmunoassay (SRL, Tokyo, Japan) and calculated urinary AVP excretion for 24h described previously (24, 25). Plasma irisin was measured using an enzyme immunoassay kit (Phoenix Pharmaceuticals Inc, Burlingame, CA, USA).

### Histological analysis

After the rats were sacrificed, kidneys were excised and decapsulated. The left kidney was immediately frozen in liquid nitrogen and the right kidney was sliced perpendicularly to the sagittal axis at approximately 5 mm intervals. Slices from the midportion of the kidneys were fixed in 10% buffered formalin overnight, and the tissue was then embedded in paraffin. Sections (3 µm thick) were stained with hematoxylin and eosin (HE), periodic acid–Schiff (PAS), and Masson’s trichrome (MT) following standard protocols. The whole kidney area and the cyst area in the HE-stained sections were determined using ImageJ analysis software (National Institutes of Health, Bethesda, MD) (26). Glomerular injury was evaluated in PAS-stained glomeruli using the index of glomerular sclerosis (13). The percentage of interstitial fibrosis area was estimated in MT-stained tissue, except for the cyst areas, glomeruli, and blood vessels, as described previously (11, 14).

### Immunohistochemical analysis

Deparaffinized kidney sections (5 μm thick) were immunostained with antibodies against desmin (ab8470, Abcam, Cambridge, UK), Ki-67 (#418071, Nichirei Biosciences, Tokyo, Japan), p-mTOR (#293133, Santa Cruz Biotechnology, Santa Cruz, CA, USA), and p-ERK (#4376, Cell Signaling Technology, Danvers, MA, USA) according to the instructions for analyzing under a light microscope (Eclipse 80i microscope, Nikon, Tokyo, Japan). For each section, 30 randomly chosen fields were photographed using a digital color camera (DS-Fi2-U3 color camera, Nikon). Using ImageJ, the stained percentage of the target area was then estimated after selecting a glomerular area with desmin staining(13). The percentage of cells positive for Ki-67, was calculated from the total number of cells containing epithelial cysts and non-cystic tubules from each kidney section using ImageJ, as described previously (27).

### Western blot analysis

The frozen kidney of each rat was thawed, dissected into the cortex and medulla, and then homogenized in 100 mmol/L potassium buffer (pH 7.25) containing 30% glycerol, 1 mmol/L dithiothreitol, and 0.1 mmol/L phenylmethylsulfonyl fluoride (14). Protein expression and phosphorylation were examined using western blot analysis, as described previously (18). Antibodies against Raf-B (#5284; Santa Cruz), ERK (#4695; Cell Signaling Technology), p-ERK (#4376; Cell Signaling Technology), mTOR (#2983; Cell Signaling Technology), p-mTOR (#2971; Cell Signaling Technology), S6 (#2217; Cell Signaling Technology), and p-S6 (#2211; Cell Signaling Technology) were used. Secondary HRP-conjugated mouse anti-rabbit (#2357; Santa Cruz) and rabbit anti-mouse (#516102; Santa Cruz) antibodies were then used. Relative band intensities were quantified using ImageJ and normalized using β-actin (A2228; Sigma-Aldrich, St. Louis, MO, USA) as an internal standard.

### cAMP assay

The frozen kidneys were ground to a fine powder with liquid nitrogen in a stainless-steel mortar. After the liquid nitrogen had evaporated, the tissues were assayed for cAMP using an enzyme-linked immunosorbent assay kit (Enzo Life Sciences Inc., Farmingdale, NY, USA) (28). Results are expressed in pmol/mg of tissue protein.

### Statistical analysis

Data are expressed as the mean ± SEM. Statistical comparisons between the groups were performed using the two-tailed unpaired *t*-test or one-way ANOVA. All analyses were carried out using GraphPad Prism software (version 8.4; GraphPad Inc., La Jolla, CA, USA). *P*-values of <0.05 were considered statistically significant.

## Results

### General parameters and urinary parameters

PCK rats as a slow progression model of PKD and Sprague-Dawley (SD) rats as a control model, were used to assess general parameters and urinary parameters in the kidney. Bodyweight was similar between the control SD rats (Con-SD) and sedentary PCK rats (Sed-PCK) groups, but was significantly lower in the exercise PCK rats (Ex-PCK) group than in the Sed-PCK group after 10 weeks of age (*P*<0.05) (Figure 1A). There were no differences in food or water intake among the three groups (Figure 1B and 1C). Urine output was similar between the Con-SD and Sed-PCK groups, but was significantly lower in the Ex-PCK group than in the Sed-PCK group at the end of the experiment (*P*<0.05) (Figure 1D). Urinary protein and liver-type fatty acid-binding protein (L-FABP) excretions were significantly increased in the Sed-PCK group after 14 weeks of age compared with the beginning of the experiment, and were significantly higher in the Sed-PCK group than in the Ex-PCK group by the end of the experiment (*P*<0.01 and *P*<0.01, respectively) (Figure 1E and 1F).

**Figure 1.**
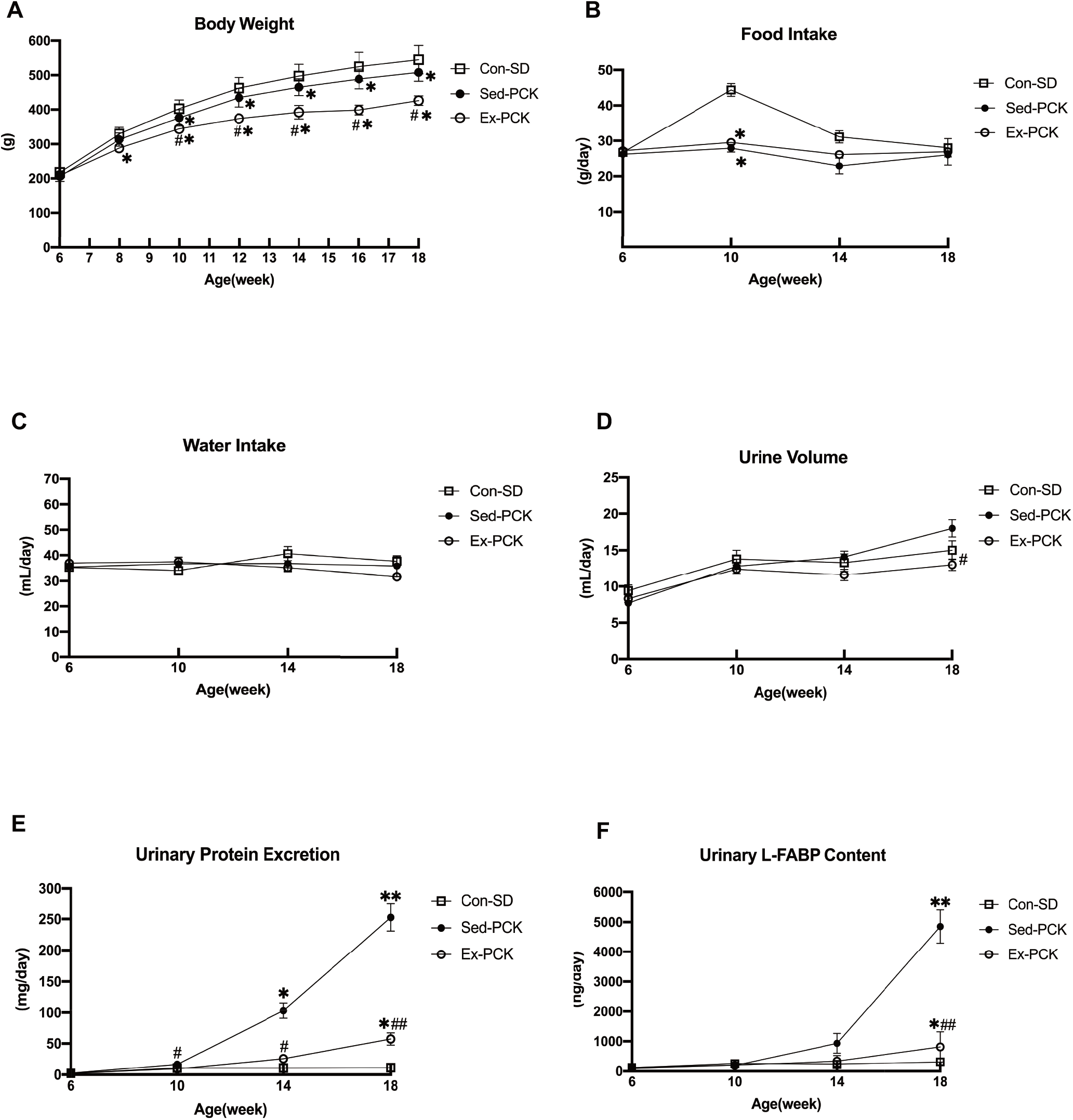
Effects of chronic exercise on general parameters and urinary parameters in PCK rats. Time courses of **(A)** body weight, **(B)** food intake, **(C)** water intake, **(D)** urine volume, **(E)** urinary protein excretion, and **(F)** urinary L-FABP excretion were compared among the Con-SD (rectangle dots), Sed-PCK (closed dots), and Ex-PCK (round dots) groups (*n*=10 in each group). Data are presented as the mean ± SEM. \**P*<0.05, **P<0.01 compared with the Con-SD group; #*P*<0.05, ##*P*<0.01 compared with the Sed-PCK group.

### Plasma parameters

Table 1 shows the plasma parameters of the groups. Total cholesterol and creatinine were significantly higher in the Sed-PCK group than in the Con-SD group, and plasma glucose was significantly lower in the Sed-PCK group than in the Con-SD group. Glucose, total cholesterol, triglyceride, urea nitrogen, and creatinine were significantly lower in the Ex-PCK group than in the Sed-PCK group. Plasma irisin was similar between the Con-SD and Sed-PCK groups, but was significantly higher in the Ex-PCK group than in the Sed-PCK or Con-SD group (*P*<0.01 and *P*<0.05, respectively).

**Table 1.** Blood pressure and plasma parameters

|  | Con-SD | Sed-PCK | Ex-PCK |
| --- | --- | --- | --- |
| Systolic Blood Pressure (mmHg) | 106 ± 7 | 104 ± 3 | 96 ± 4 |
| Glucose (mg/dL) | 169.1 ± 6.8 | 139.7 ± 7.8* | 125.5 ± 20.1* |
| Total cholesterol (mg/dL) | 67.9 ± 5.1 | 148.2 ± 8.6* | 114.1 ± 7.6*# |
| Triglyceride (mg/dL) | 59.7 ± 6.0 | 69.3 ± 5.3 | 49.8 ± 3.7# |
| Urea nitrogen (mg/dL) | 16.8 ± 0.5 | 18.2 ± 0.9 | 15.9 ± 0.3# |
| Creatinine (mg/dL) | 0.28 ± 0.01 | 0.35 ± 0.02* | 0.30 ± 0.01# |
| Irisin (ng/dL) | 1134.8 ± 29.7 | 1070.0 ± 41.0 | 1578.1 ± 106.0*## |

Con-SD, control Sprague-Dawley rats; Sed-PCK, sedentary polycystic kidney rats; Ex-PCK, exercise polycystic kidney rats. Data are presented as means±SEM. *n*=10 in each group. \**P*<0.05 compared with the Con-SD group; #*P*<0.05, ##*P*<0.01 compared with the Sed-PCK group.

### Kidney weight and morphology

Figure 2A shows representative images of the HE-stained kidney from the three groups. Renal cysts were observed in the outer medulla of both the Sed-PCK and Ex-PCK groups, and cyst sizes were smaller in the Ex-PCK group than in the Sed-PCK group. Total kidney weight was significantly lower in the Ex-PCK group than in the Sed-PCK group (*P*<0.01) (Figure 2B), but the kidney-to-body weight ratio was not significantly different between the two PCK groups (Figure 2C). The cystic index was significantly higher in the Sed-PCK group than in the Con-SD group (*P*<0.01), and significantly lower in the Ex-PCK group than in the Sed-PCK group (*P*<0.05) (Figure 2D).

**Figure 2.**
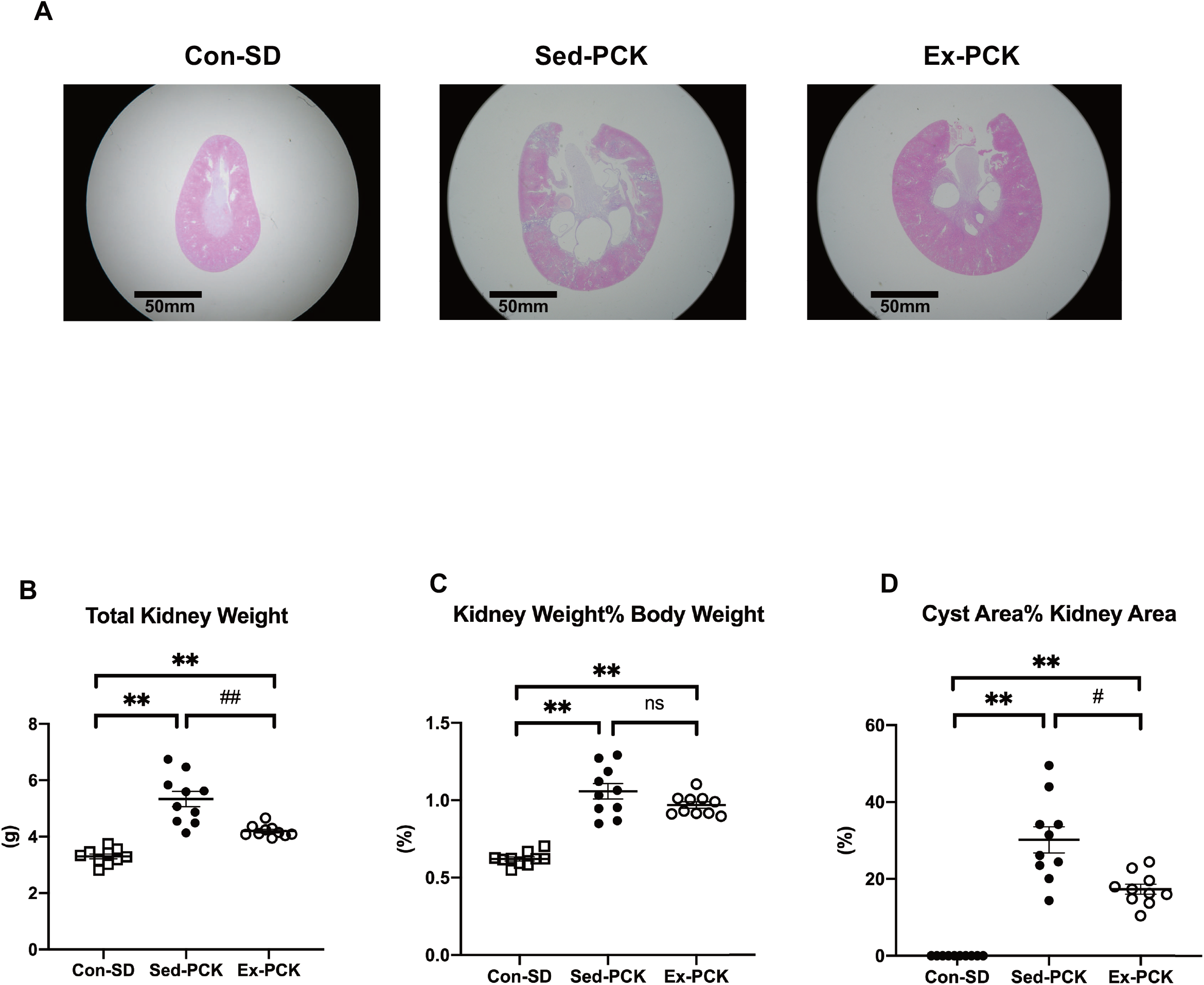
Effects of chronic exercise on kidney cysts in PCK rats. **(A)** Representative images of kidney specimens stained with HE in the Con-SD, Sed-PCK, and Ex-PCK groups. **(B)** Total kidney weight, **(C)** kidney-to-body weight ratio, and **(D)** cystic index were compared among the Con-SD (rectangle dots), Sed-PCK (closed dots), and Ex-PCK (round dots) groups (*n*=10 in each group). Data are presented as the mean ± SEM. \*\**P*<0.01 compared with the Con-SD group; #*P*<0.05 compared with the Sed-PCK group; ns: no significant difference.

### Glomerular damage and renal interstitial fibrosis

Figure 3A shows representative images of PAS-stained and desmin-immunostained glomeruli and MT-stained kidneys in each group. Glomerular sclerosis, podocyte injury, and renal interstitial fibrosis were observed in the Sed-PCK group. The index of glomerular sclerosis was significantly higher in the Sed-PCK group than in the Con-SD group (*P*<0.01), and significantly lower in the Ex-PCK group than in the Sed-PCK group (*P*<0.05) (Figure 3B). The desmin-positive staining area in the glomeruli was significantly larger in the Sed-PCK group than in the Con-SD group (*P*<0.01), and significantly smaller in the Ex-PCK group than in the Sed-PCK group (*P*<0.01) (Figure 3C). The renal interstitial fibrosis area was significantly higher in the Sed-PCK group than in the Con-SD group (*P*<0.01), and smaller in the Ex-PCK group than in the Sed-PCK group (Figure 3D).

**Figure 3.**
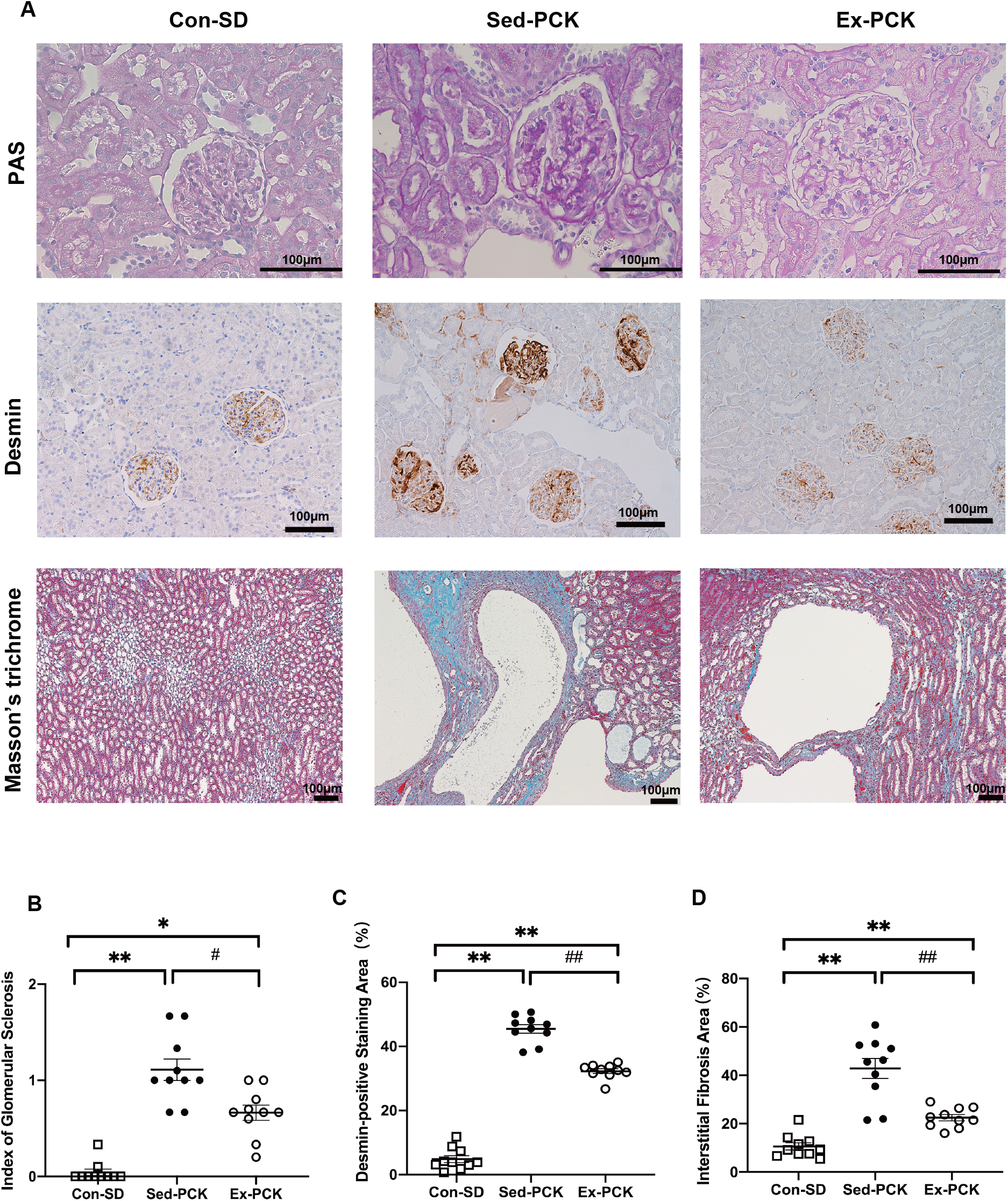
Effects of chronic exercise on glomerular sclerosis, podocyte injury, and renal interstitial fibrosis in PCK rats. **(A)** Representative images of periodic acid–Schiff (PAS)-stained, desmin-immunostained glomeruli and Masson’s trichrome stained kidneys in the Con-SD, Sed-PCK, and Ex-PCK groups. **(B)** Index of glomerular sclerosis, **(C)** desmin-positive staining area (%), and **(D)** interstitial fibrosis area (%) in the Con-SD (rectangle dots), Sed-PCK (closed dots) and Ex-PCK (round dots) groups (*n*=10 in each group). Data are presented as the mean ± SEM. \**P*<0.05, \*\**P*<0.01 compared with the Con-SD group; #*P*<0.05, ##*P*<0.01 compared with the Sed-PCK group.

### Cell proliferation and signaling cascades

Figure 4A shows representative images of the kidney immunostained for Ki-67 from the Sed-PCK and Ex-PCK groups. Ki-67-positive cells were highly expressed in the cyst-lining epithelium, interstitium, and non-cystic tubules of the Sed-PCK group. Chronic exercise led to fewer Ki-67-positive cells. The Ki-67 labeling index was significantly lower in the cyst-lining epithelium and non-cystic tubules in the Ex-PCK group compared with the Sed-PCK group (*P*<0.01 and *P*<0.01, respectively) (Figure 4B and 4C).

**Figure 4.**
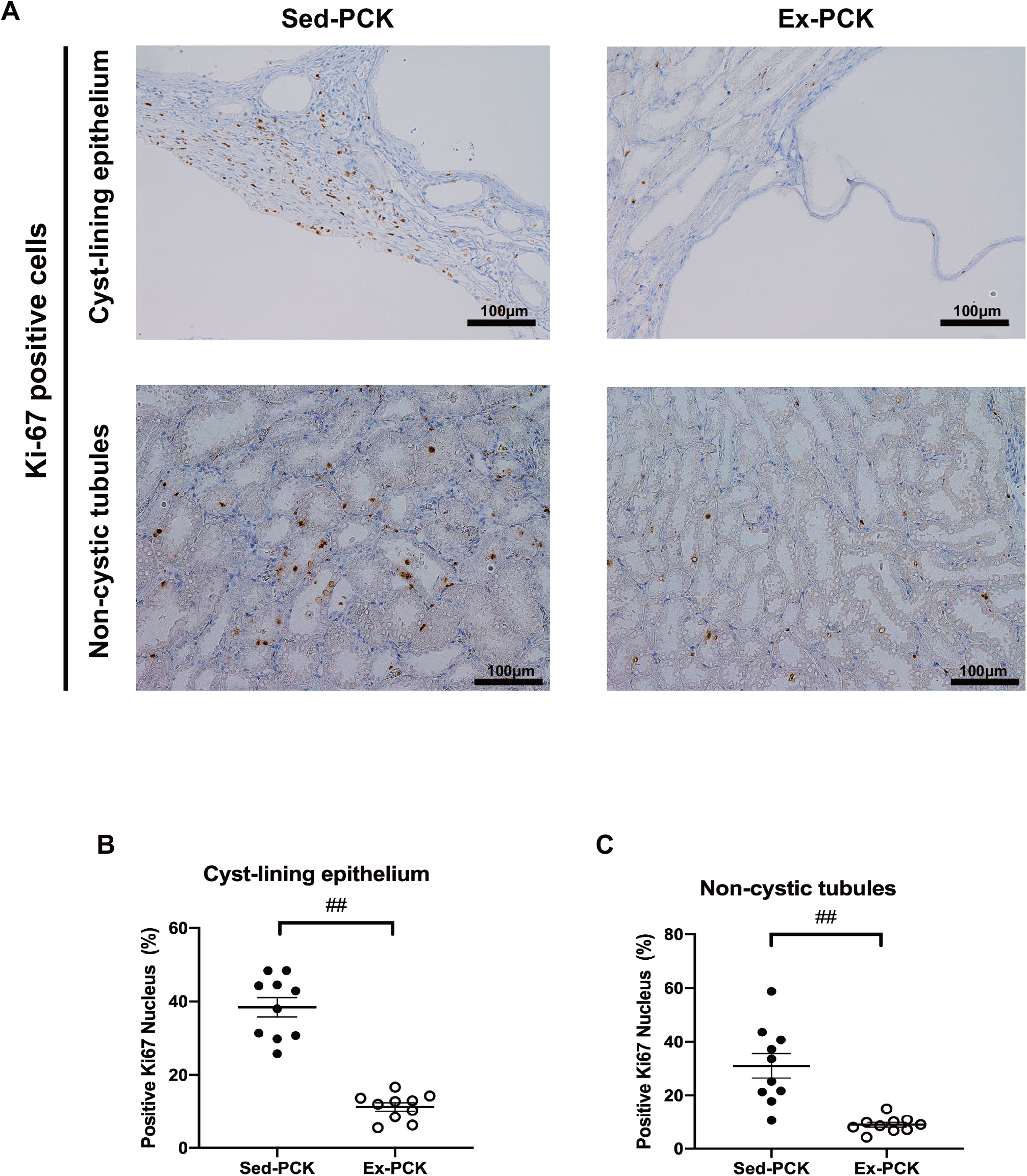
Effects of chronic exercise on cell proliferation in the kidneys of PCK rats. **(A)** Representative images of kidney specimens immunostained for Ki-67 in the Sed-PCK and Ex-PCK groups. **(B)** Ki-67 labeling index in the cyst-lining epithelium of the Sed-PCK (closed dots) and Ex-PCK (round dots) groups (*n*=10 in each group). Ki-67 labeling index in the non-cystic tubules of the Sed-PCK (closed dots) and Ex-PCK (round dots) groups (*n*=10 in each group). Data are presented as the mean ± SEM. ##*P*<0.01 compared with the Sed-PCK group.

Urinary AVP excretion was significantly higher in the Sed-PCK group than in the Con-SD group (*P*<0.05), and was considerably higher in the Ex-PCK group than in the Sed-PCK group (*P*<0.05) (Figure 5A). Renal cAMP content was significantly higher in the Sed-PCK group than in the Con-\ SD group (*P*<0.05), but it was not significantly different between the Sed-PCK and Ex-PCK groups (Figure 5B). Renal B-Raf expression was significantly higher in the Sed-PCK group than in the Con-SD (*P*<0.01), and significantly lower in the Ex-PCK group than in the Sed-PCK group (*P*<0.01) (Figure 5C).

**Figure 5.**
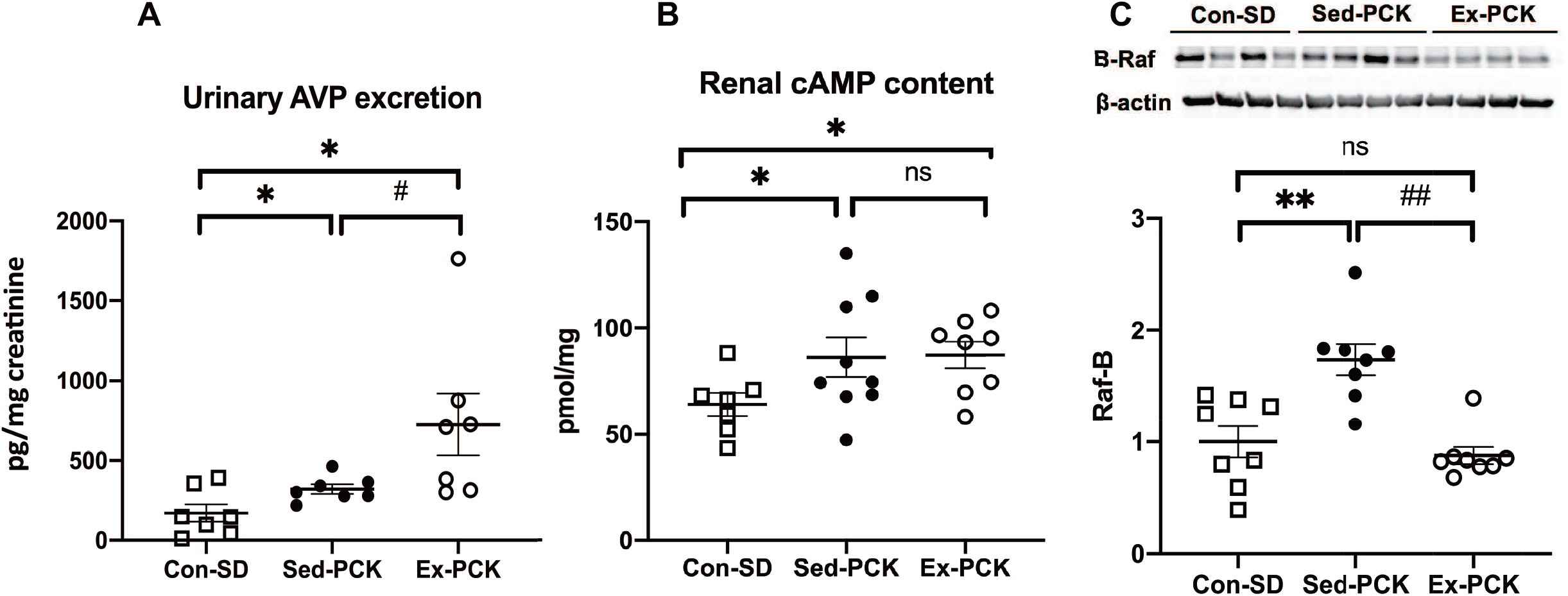
Effects of chronic exercise on urinary AVP excretion, renal cAMP content, and renal B-Raf expression in PCK rats. **(A) U**rinary AVP excretion in the Con-SD (rectangle dots), Sed-PCK (closed dots), and Ex-PCK (round dots) groups. **(B)** Renal cAMP content in the Con-SD (rectangle dots), Sed-PCK (closed dots), and Ex-PCK (round dots) groups (*n*=10 in each group). **(C)** Western blotting analysis of B-Raf expression in the Con-SD (rectangle dots), Sed-PCK (closed dots), and Ex-PCK (round dots) groups (*n*=8 in each group). Top panels show representative immunoblotting. Each lane was loaded with a protein sample prepared from four different rats per group. The ratio in the Con-SD group was assigned a value of 1. Data are presented as the mean ± SEM. \**P*<0.05, \*\**P*<0.01 compared with the Con-SD group; #*P*<0.05, ##*P*<0.01 compared with the Sed-PCK group; ns: no significant difference.

Figure 6A and 6B show representative images of kidneys immunostained for phosphorylated (p-) ERK and p-mTOR, respectively, from each group. The p-ERK and p-mTOR proteins were highly expressed in the cyst-lining epithelium and non-cystic tubules in the Sed-PCK group, and chronic exercise decreased their expressions (Figure 6A and 6B). Renal ERK and mTOR phosphorylation were significantly higher in the Sed-PCK group than in the Con-SD group (*P*<0.01 and *P*<0.01, respectively), and S6 phosphorylation also tended to be higher in the Sed-PCK group compared with the Con-SD group (Figure 6C, 6D, and 6E). Renal ERK, mTOR, and S6 phosphorylation was significantly lower in the Ex-PCK group than in the Sed-PCK group (*P*<0.01, *P*<0.01, and *P*<0.01, respectively).

**Figure 6.**
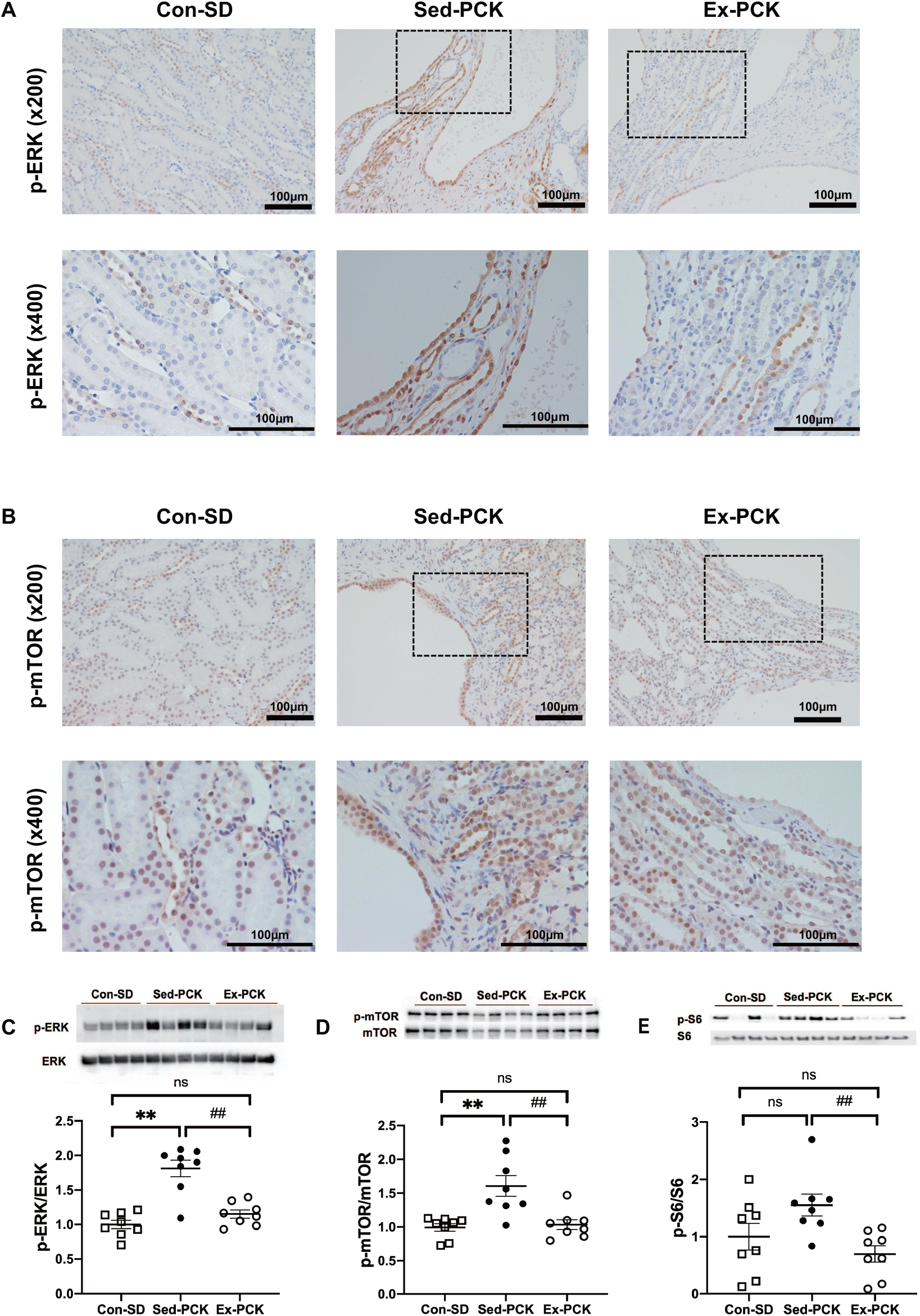
Effects of chronic exercise on the phosphorylation of ERK, mTOR, and S6 in PCK rats. Representative images of kidney specimens immunostained for **(A)** p-ERK and **(B)** p-mTOR in the Con-SD, Sed-PCK, and Ex-PCK groups. Western blotting analysis of **(C)** p-ERK, **(D)** p-mTOR, and **(E)** p-S6 expression in the Con-SD (rectangle dots), Sed-PCK (closed dots), and Ex-PCK (round dots) groups (*n*=8 in each group). Top panels show representative immunoblotting. Each lane was loaded with a protein sample prepared from four different rats per group. Ratios of the relative band intensity of the phosphorylated protein to that of the total protein were calculated. The ratio in the Con-SD group was assigned a value of 1. Data are presented as the mean ± SEM. \*\**P*<0.01 compared with the Con-SD group; ##*P*<0.01 compared with the Sed-PCK group; ns: no significant difference.

## Discussion

Chronic exercise has renal protective effects in CKD patients and models (10-14); however, the renal protective effects of chronic exercise have not yet been reported in PKD patients or models. The present study revealed that chronic exercise at a moderate intensity slowed the progression of renal cyst growth, glomerular damage, interstitial fibrosis, and renal dysfunction in PCK rats, despite increasing AVP. Chronic exercise also inhibited excessive cell proliferation, with down-regulation of the cAMP/B-Raf/ERK and mTOR/S6 pathways in renal epithelial cells. To the best of our knowledge, the present study is the first to report that chronic exercise has therapeutic potential against cyst growth and renal dysfunction in PKD.

We chose the exercise protocol in the present study based on our previous study of CKD model rats with 5/6 nephrectomy (11), in which proteinuria and glomerular sclerosis were significantly attenuated after 12 weeks of chronic exercise. We confirmed that when PCK rats run at a speed of 28 m/min on the treadmill, oxygen consumption (VO_2_) corresponds to approximately 65% of the maximal VO_2_, which is assumed to be aerobic exercise at a moderate intensity (18). In contrast to the present results, Darnley et al. reported that treadmill exercise (14 m/min, 30 min/day, 3 days/week) for 6 weeks did not lead to any changes in serum urea nitrogen or creatinine in Han:SPRD-*cy* rats (19). Similarly, in our pilot studies, chronic exercise for 8 weeks did not significantly affect renal cyst growth in PCK rats (data not shown). Thus, the intensity, time, frequency, and duration of the exercise protocol may be important to obtain benefits in PKD models. In agreement with our previous studies (10-13), chronic exercise lowered proteinuria and plasma creatinine and attenuated glomerular sclerosis and podocyte injury in PCK rats. Chronic exercise for 8 weeks significantly decreased urinary protein excretion (Figure 1E) without significant effects on renal cyst growth in PCK rats (data not shown). Therefore, glomerular protection may be a primary effect of chronic exercise, rather than being secondary to slowing renal cyst growth. Urinary L-FABP excretion, a biomarker of proximal tubular stress and tubulointerstitial disorder, increases linearly with age and reflects the progression of tubulointerstitial disorder in PCK rats (29). Chronic exercise might therefore strongly attenuate proximal tubular stress and tubulointerstitial disorder in PCK rats. As an indicator of cell proliferation, chronic exercise decreased the number of Ki-67-positive cells in the kidneys of PCK rats, indicating the inhibition of excessive cell proliferation. As well as in the kidney, chronic exercise has also been recently reported to slow the progression of cyst growth and fibrosis in the liver of PCK rats (18).

The present study indicates that chronic exercise increases AVP in PCK rats. In agreement with these results, AVP synthesis and secretion have been previously reported to increase during exercise (30). Sustained moderate exercise (at an intensity threshold of 40%–65% of VO_2max_) increased plasma AVP (31, 32). Furthermore, chronic exercise with a treadmill for 5 weeks increased plasma AVP in Wistar rats (33). The present study also indicates that chronic exercise did not change renal cAMP content and did decrease the cAMP-inducible B-Raf expression in PCK rats, despite increasing AVP, suggesting that chronic exercise might inactivate adenylate cyclase via the inhibitory G protein (Gi). Previous studies indicate that norepinephrine and α2-adrenergic receptor (α2-AR) agonists inhibit the AVP-activated adenylate cyclase, cAMP content, and water transport in the collecting ducts (34-36). Therefore, it is possible that chronic exercise might stimulate renal sympathetic activity and activate α2-AR in the collecting ducts to slow the progression of renal cyst growth with reducing the renal cAMP content in PCK rats. In this regard, our preliminary study indicates that chronic treatment of the α2-AR agonist, clonidine slows the progression of renal cyst growth in PCK rats (data not shown).

Previous studies indicate that even normal plasma AVP levels increase B-Raf expression and ERK phosphorylation in the kidneys of PCK rats, and that inhibition of AVP by V2R antagonists and hydration can down-regulate the B-Raf/ERK pathway (24, 37). The present study indicates that chronic exercise down-regulates not only the B-Raf/ERK pathway but also the mTOR/S6 pathway in the kidneys of PCK rats. Both mTOR and ERK are involved in excessive cell proliferation and cyst growth in the renal tubules and cholangiocytes of PCK rats (20). However, neither tolvaptan nor an ERK inhibitor, AEZ-131, affected S6 phosphorylation in the kidney of PCK rats, and the suppressive effects of tolvaptan and an mTOR inhibitor, rapamycin, on renal cyst growth were additive (38). The suppressive effects of chronic exercise on excessive cell proliferation and renal cyst growth in the present study might therefore be mediated by down-regulation of both the B-Raf/ERK and mTOR/S6 pathways in the kidneys of PCK rats. Several types of exercise affect mTOR and ERK in the skeletal muscle, fat, liver and vasculature (39-41). However, the effects of exercise on mTOR or ERK have not previously been reported in the kidney, especially in the renal tubules. We recently reported that chronic exercise down-regulates mTOR and ERK phosphorylation in the liver and cholangiocytes in PCK rats (18). In agreement with the results from PCK rats, chronic exercise with a treadmill inactivated mTOR and suppressed excessive cell proliferation in hepatocellular carcinoma in PTEN-deficient mice (42) and carcinoma-implanted rats (43).

The present study also demonstrates that chronic exercise increases plasma irisin in PCK rats. Irisin mediates the beneficial effects of exercise, such as by promoting the brown adipose formation and improving the metabolism, and also has a beneficial role in kidney and heart diseases (15, 44-46) In one study, plasma irisin levels were significantly decreased in CKD patients and were inversely correlated with blood urea nitrogen and creatinine levels (47). In another study, skeletal muscle-specific PGC-1α overexpression increased irisin production and plasma irisin levels and attenuated renal damage in mice with folic acid nephropathy, unilateral ureteral obstruction, and 5/6 nephrectomy (15). Moreover, recombinant irisin administration attenuated renal damage in the mouse kidney disease models (15). Although it is unknown whether irisin can inhibit cyst growth, irisin has been reported to inhibit mTOR, ERK, and cell proliferation in cultured cardiomyocytes, cardiomyoblasts, and pancreatic cancer cells (48, 49). Additionally, irisin increased intracellular Ca^2+^ in cultured cardiomyoblasts and endothelial cells (50). Future study is necessary to examine whether irisin directly acts on renal epithelial cells and inhibits cyst growth in PCK rats.

In conclusion, chronic exercise slows the progression of PKD pathologies, such as renal dysfunction, renal cyst growth, glomerular damage, and renal interstitial fibrosis in PCK rats. Despite increasing AVP, chronic exercise also inhibits excessive cell proliferation, with down-regulation of the cAMP/B-Raf/ERK and mTOR/S6 pathways in the kidney of PCK rats. Although the results of the present study may not be directly applicable to humans, chronic exercise may be a novel therapeutic approach against cyst growth and renal dysfunction in PKD patients.

## Acknowledgments

This work was supported by Grants-in-Aid for Scientific Research from Japan Society for the Promotion of Science grants 15K12573, 17H02119, 20H04054, 20J12732 and 20K19338. We greatly appreciate the technical support received from the Biomedical Research Unit of Tohoku University Hospital, Animal Pathology Platform and Biomedical Research Core of Tohoku University Graduate School of Medicine for the histopathological analysis. The results of the study are presented clearly, honestly, and without fabrication or inappropriate data manipulation. The results of the present study do not constitute endorsement by the American College of Sports Medicine.

## Conflict of Interest

The authors declare no conflicts of interest associated with this manuscript.

## Notes

### Competing Interest Statement

The authors have declared no competing interest.

